# An insight into cellulolytic capacity of the *Trichoderma harzianum* P49P11 revealed by omics approaches

**DOI:** 10.1101/2022.06.19.496725

**Authors:** Carla Aloia Codima, Geizecler Tomazetto, Gabriela Felix Persinoti, Diego M. Riano-Pachon, Fábio Marcio Squina, José Geraldo da Cruz Pradella, Priscila da Silva Delabona

**Author notes:** Corresponding author: Geizecler Tomazetto. These authors contributed equally to this work (Carla Aloia Codima, Geizecler Tomazetto).

## Abstract

Cellulases are a group of enzymes with several applications in biofuel production, and the paper, food, pharmaceutical, and chemical industries. *Trichoderma harzianum* P49P11 secrete all cellulases with high efficiency, representing an alternative to the current filamentous fungi in biotechnological industries. In this study, the cellulolytic mechanisms employed by the strain P49P11 to degrade crystalline cellulose in batch fermentation culture mode were elucidated by combining genome and secretome analysis. The strain P49P11 encodes nineteen cellulase genes from five different CAZyme families (GH5, GH6, GH7, GH12, and GH45), followed by several enzyme families for hemicellulose, pectin, and alpha-and beta-glucans degradation. The diverse CAZymes were also observed in the secretome, including cellulases, hemicellulases, and glucanases. In addition, β-glucosidases and xylanase activities detected during the fermentation process validated our secretome analysis. Taken together, our results revealed all enzymatic machinery used by the *T. harzianum* P49P11 to degrade cellulose in batch fermentation mode.

**Highlights:**

- We described a high-quality genome assembly and annotation of the *T. harzianum* P49P11.
- The *T. harzianum* P49P11 genome possesses a complete set of genes for lignocellulose degradation.
- The first report on *T. harzianum* P49P11 secretome obtained from batch fermentation strategy.
- *T. harzianum* P49P11 produced cellulases, lignocellulases, and auxiliary enzymes produced in response to crystalline cellulose.

## Background

Cellulases refer to a group of enzymes consisting of three different classes of enzymes (endoglucanases, exoglucanases, and β-glucosidases) required to completely degrade cellulose. Endoglucanases cleave glycosidic bonds to produce reducing and non-reducing ends of cellulose chains at which exoglucanases can act to release some glucose and cellobioses, then β-glucosidases hydrolyse cellobiose into glucose completing the saccharification process. Furthermore, cellulases are classified in several glycosyl hydrolases (GH) families as defined by the CAZy database [1], including GH5, GH7, GH12, and GH45.

Cellulases have several industrial applications (e.g., in the paper, food, pharmaceutical, and chemical industries) representing 10% of the enzymes market valued at USD 10.7 billion in 2020, and this market is expected to grow to USD 17.88 billion by 2027 [2]. However, the most significant application lies in commercial enzymatic cocktails used in lignocellulose-based biorefineries [3][4][5].

Fungi are well-known for their capacity to secrete a broad range of cellulases with distinct proprieties and high titers – which are desired for the large application and demand for biotechnological purposes [3][4] [6]. *Trichoderma reesei* is the main filamentous fungi (hereafter called fungi) widely used to produce cellulases in industry owing to the high capacity of secretion – reaching up 100 g/l of cellulase [7][6] [5]. However, although *T. reesei* produce substantial amounts of exoglucanases and endoglucases, β-glucosidases production represents a low percentage of the total cellulases secreted, leading an incomplete saccharification process [8]. Therefore, due to the variety of applications and market expansion, there is still a need to identify and explore novel fungi capable of producing a high level of different cellulases at the same time with good activity and stability for efficiently digesting cellulose into glucose [4][6].

Some *T. harzianum* strains have a high capacity to produce a large amount of all cellulases – including with high β-glucosidases activity [9][10][11] [12] [13], representing a potential alternative to *T. reesei* for cellulase production on an industrial scale. Examples include the utilization of *T. harzianum* P49P11 for production of cellulases in a stirred tank bioreactor using bagasse as a carbon source [9] [10], or in continuous fermentation process [14]. Furthermore, the crude extract produced by strain P49P11 has shown similar lignocellulolytic performance as produced by *T. reesei* RUTC30, indicating the strain P49P11 as a potential alternative producer for on-site production of enzymes [15]. Therefore, studies using omics approaches are still required to elucidate the genetic content and identify the cellulases genes expressed by strain P49P11 under different conditions, improving our understanding of its microbial physiology and cellulolytic potential.

In this context, the cellulolytic mechanisms of *T. harzianum* P49P11 were investigated in a stirred tank bioreactor using omics approaches. For this purpose, we first sequenced the P48P11 strain genome to reveal all its CAZy families. To link the genetic potential with enzyme secretion capacity of the P49P11, its secretome in response to microcrystalline cellulose was analyzed using LC-MS/MS. In addition, we performed a time course analysis of the enzyme activity profiles over 5 days of the fermentation process. Taken together, our findings provide a specific description of how cellulose is degraded by P49P11 in a batch bioreactor experiment.

## Results and discussion

### Genome sequence

The assembled P49P11 genome was 42.05 Mb, distributed across 919 contigs with N50 of 181.635 and GC content of 48.6%. Table 1 summarizes the genomic features of strain P49P11 genome. Gene prediction identified 11,755 genes, and among them 81.88% and 51% exhibit similarity to known function annotated proteins in Pfam, KOG (Eukaryotic orthologous groups – a eukaryote-specific version of the COGs) databases. The complete annotation is available in Supplementary Material S1. The completeness and integrity of the draft genome assembly, determined by Benchmarking Universal Single-Copy Orthologs (BUSCO) analysis, revealed 98.3 % BUSCO genes and 0.3 % fragmented BUSCOs (Table S2).

**Table 1.** Genome assembly and annotation statistics of the *Trichoderma harzianum* P49P11

| Assembly statistics | *T. harzianum* P9P11 |
| --- | --- |
| Estimated coverage | 250 |
| Genome size (Mbp) | 42.05 |
| Contigs ($\geq 0$ bp) | 919 |
| Contigs ($\geq 1000$ bp) | 919 |
| Contigs ($\geq 5000$ bp) | 382 |
| Contigs ($\geq 10\,000$ bp) | 233 |
| Contigs ($\geq 25\,000$ bp) | 129 |
| Contigs ($\geq 50\,000$bp) | 102 |
| Largest Contig | 1686451 |
| N50 | 497678 |
| N75 | 232825 |
| L50 | 26 |
| L75 | 56 |
| % GC | 48.6 |
| Number of protein coding genes | 11,755 |
| Proteins with predicted Pfam domain | 9,626 |
| Proteins assigned to KOG | 5,996 |

### *CAZyme profile of T. harzianum* P49P11

The *T. harzianum* P49P11 genome encodes 450 protein sequences with at least one CAZyme domain, including 259 glycoside hydrolase (GHs) modules, 23 carbohydrate esterase (CE), 13 carbohydrate-binding modules (CBMs), 72 auxiliary activity (AA), 11 polysaccharide lyase (PL), and 85 glycosyltransferases (GT) (Fig.1 and Supplementary Material S2).

**Figure 1.**
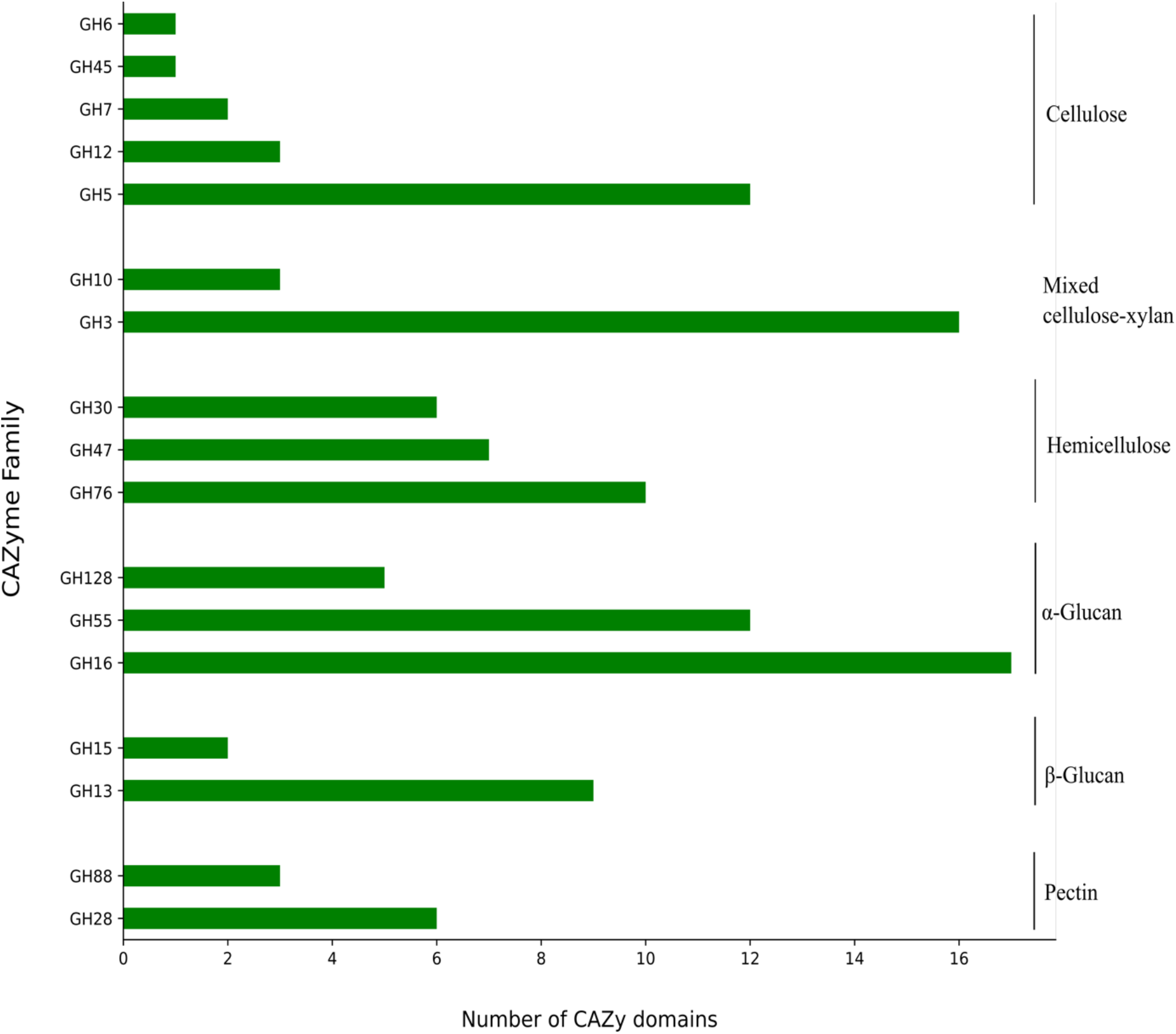
CAZyme profile of the *Trichoderma harzianum* P49P11 revealed by genomic analysis. The most abundant GH families in the P49P11 genome involved in lignocellulose degradation are displayed.

Analyzing in more detail the CAZyme profile, strain P49P11 genome encodes GH families associated with cellulose (e.g., GH5, GH12, GH45), hemicellulose (GH76, GH47, GH30), alpha-glucan (GH13, GH15), beta-glucan (GH16, GH55), and chitin (GH18) degradation. The strain encodes all enzymes for complete cellulose deconstruction, including endoglucanases, exoglucanases, β-glucosidases from GH5 (12 genes), GH6 (1), GH7(2), GH12 (3), and GH45 (1), totalizing 19 genes. However, the most abundant genes encoding GH families were associated with chitin degradation (GH18, 27 genes). Although *Trichoderma* species are well-known as a power producer of cellulases [16][17], their genomes have been described with a higher number of GH18 [18]. The abundance of genes encoding chitinolytic enzymes might be explained by the mycoparasitism function of some *Trichoderma* species, which produce such enzymes to attack chitin from fungal cells [19].

Of the predicted CE families, eight different families predicted were mostly represented by CE5 family (6 genes), which are related to cutinase and acetyl xylan esterase activities. Eleven different AA families were predicted, predominantly represented by AA3_2 (17 genes) and AA3_3 (two genes), of which some are classified as oxidases – enzymes act on crystalline cellulose and on lignin linkages [20][1]. Furthermore, strain P49P11 genome also possesses 3 copies of lytic polysaccharide monooxygenases-encoding genes (LPMOs – AA9), which catalyzes oxidative cleavage of cellulose and other polysaccharides [20]. Four PLs were families detected, with PL7 alginate lyases as the predominant ones. Therefore, *T. harzianum* P49P11 is capable to encode all enzymes require degrading lignocellulosic plant material.

For comparative purposes, we carried out the CAZymes prediction in four genomes of the *T. harzianum* strains (T6776, TR1, TR274, and CBS226) using the same criteria as those used previously for P49P11 (Supplementary Material S3). The number of genes encoding CAZymes and the distribution of the families are highly similar among the strains. These findings suggest that the difference in lignocellulolytic capacity among the *T. harzianum* strains might be much more related with regulation of gene expression than CAZyme numbers present or with dissimilarity among the sequences [21].

### Enzyme activities induced by crystalline cellulose

To evaluate the cellulase production of *T. harzianum* P49P11, the strain was grown in MA medium with 2% crystalline cellulose as the sole carbon source in batch fermentation. Besides determined cellulase production by FPase activity, we also measured β-glycosidase and xylanase activities. The presence of β-glycosidase activity suggests the complete cellulose saccharification process. The secretion of xylanases might be induced by a certain amount of xylan associated with cellulose [22][23].

PFase activity reached maximum values at 120 h (0.88 FPU m.L^-1^) of fermentation (Figure 2A). However, the high amounts of extracellular proteins reached between 72 h and 96 h with no significant differences. Similarly, the highest activities of β-glycosidase and xylanase activities were reached at 72 h of cultivation, 2.21 U / mL and 220 U / mL, respectively (Figure 2B). As the productivity (FPU L^-1^ h^-1^) achieved maximum values (10.14 FPU L^-1^ h^-1^) and extracellular protein concentration at 72 h, the secretome analysis was carried out at this time point.

**Figure 2.**
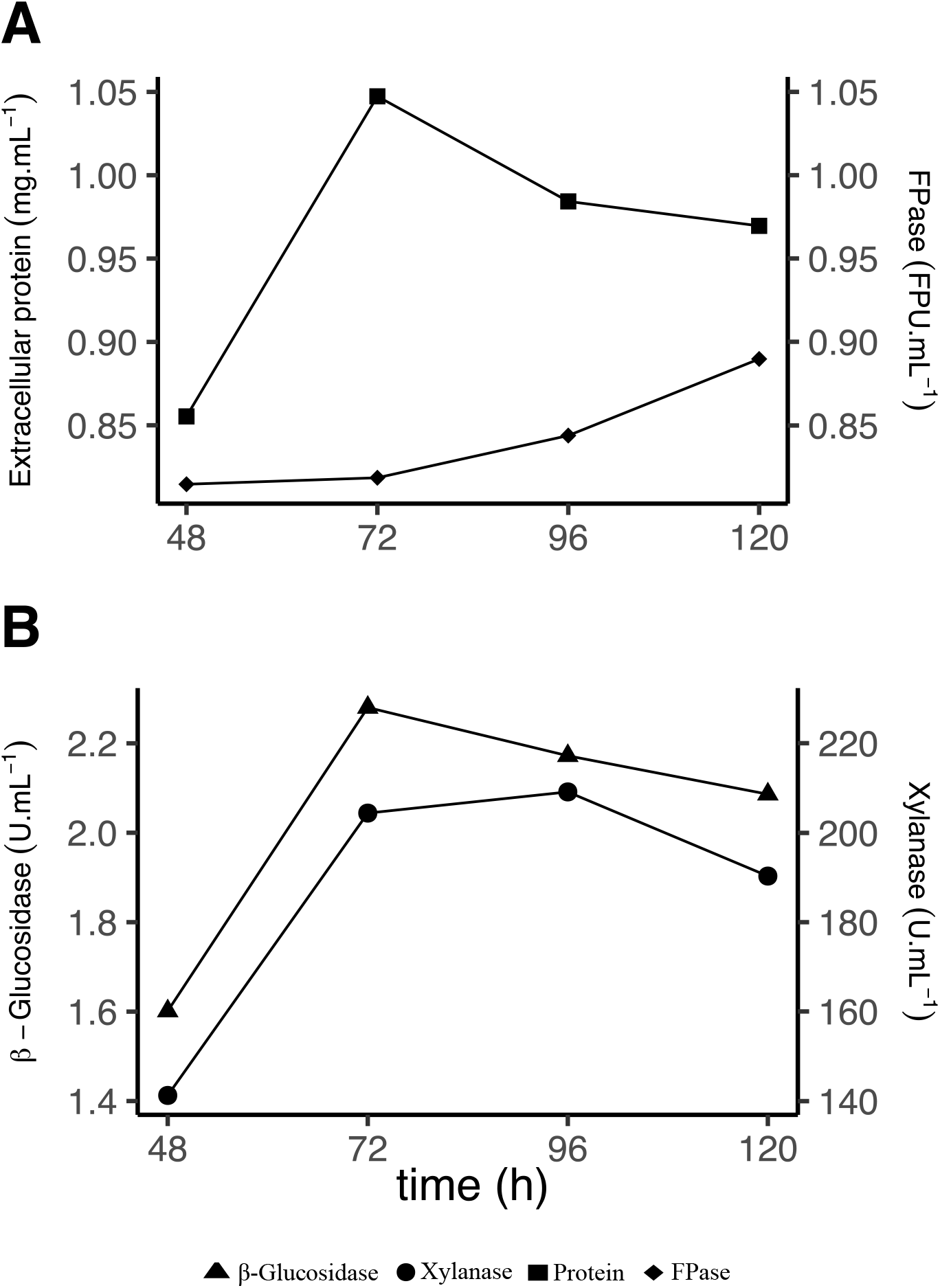
Enzymatic activities and extracellular protein production by the *Trichoderma harzianum* P49P11 cultivated in crystalline cellulose. *T. harzianum* P49P11 was grown on 2 % (w/v) of crystalline cellulose as the sole carbon source at the bench-scale in submerged fermentation. **A** Beta-glucosidase and xylanase activities measured from supernatant samples of P49P11. **B**. Extracellular protein production and PFase measured from supernatant samples of P49P11.

### *Secretome profile of T. harzianum* P49P11 *on crystalline cellulose*

Based on a minimum of two unique peptides per protein identified, 59 proteins were detected in the secretome of *T. harzianum* P49P11 cultivated in crystalline cellulose. Among them were 32 GHs from 23 different CAZymes families, followed by 4 CBMs, 2 CEs, and 2 AAs (Table 2 and Supplementary Material S3). All these CAZyme sequences were manually analyzed for the presence of extra domains that are not automatically annotated. Furthermore, most of these CAZymes were predicted with a signal peptide, thus indicating secretion of these enzymes. Hemicellulases were the most abundant enzymes in terms of CAZymes numbers detected. However, cellulases detected displayed a high spectrum count, followed by the mixed cellulase-xylanases group, accounting for 25% and 19% of the total spectra, respectively. Among 19 genes encode cellulases by strain P49P11, six sequences were detected, including one GH6 and GH12, two GH5 and GH7, which are families encoding endoglucanases, exoglucanases, and β-glucosidases. The GH7 cellulase with CBM1 appended represents the enzymes with the most spectrum detected. It is worth mentioning that CBM1 are modules exclusively found in fungi that specifically bind to crystalline cellulose. Four of six cellulases detected were fused to CBM1. In addition, a protein sequence with CBM1 appended without the catalytic domain detected might represent a novel cellulase domain. Therefore, FPase and β-glycosidase activities previously described might be a result of secretion of these cellulases families.

**Table 2.** CAZymes identified by LC-MS/MS in the secretome of the *Trichoderma harzianum* P49P11 grown in crystalline cellulose.

| Target Substrate | Protein ID | CAZy family<sup>1</sup> | Molecular Weight | Signal P<sup>2</sup> | Unique peptide<sup>3</sup> | Spectrum count<sup>4</sup> |
| --- | --- | --- | --- | --- | --- | --- |
| Cellulose | THAR_004903_T0 | GH7 - CBM1 | 53 kDa | Yes | 19 | 316 |
|  | THAR_010035_T0 | GH5 - CBM1 | 45 kDa | Yes | 4 | 10 |
|  | THAR_007673_T0 | GH5_7 | 100 kDa | Yes | 3 | 4 |
|  | THAR_010517_T0 | GH6 - CBM1 | 51 kDa | Yes | 10 | 38 |
|  | THAR_000270_T0 | GH7-CBM1 | 50 kDa | Yes | 5 | 15 |
|  | THAR_005887_T0 | GH12 | 29 kDa | Yes | 4 | 20 |
| Mixed Cellulose/Xylan | THAR_010018_T0 | GH3 | 78 kDa | Yes | 17 | 50 |
|  | THAR_005508_T0 | GH10 | 38 kDa | Yes | 6 | 16 |
|  | THAR_004929_T0 | GH10 | 44 kDa | No | 16 | 246 |
| Glucans<sup>1</sup> | THAR_000837_T0 | GH16 | 31 kDa | Yes | 5 | 13 |
|  | THAR_009992_T0 | GH16 | 34 kDa | Yes | 3 | 13 |
|  | THAR_007389_T0 | GH55 | 80 kDa | Yes | 8 | 13 |
|  | THAR_006343_T0 | GH55 | 80 kDa | Yes | 2 | 2 |
|  | THAR_008676_T0 | GH55 | 89 kDa | No | 12 | 27 |
| Hemicellulose | THAR_001809_T0 | GH62 | 35 kDa | Yes | 6 | 42 |
|  | THAR_011526_T0 | GH43 | 59 kDa | No | 3 | 4 |
|  | THAR_009358_T0 | GH92 | 88 kDa | Yes | 13 | 23 |
|  | THAR_004055_T0 | GH54 | 53 kDa | Yes | 6 | 15 |
|  | THAR_002305_T0 | GH115 | 122 kDa | No | 5 | 10 |
|  | THAR_000685_T0 | GH30 | 50 kDa | No | 4 | 13 |
|  | THAR_004907_T0 | GH30 | 52 kDa | Yes | 8 | 19 |
|  | THAR_004578_T0 | GH30 | 51 kDa | Yes | 5 | 11 |
|  | THAR_004421_T0 | GH30 | 52 kDa | Yes | 6 | 6 |
|  | THAR_000744_T0 | GH11 | 26 kDa | Yes | 6 | 151 |
| Oligosaccharideos | THAR_007271_T0 | GH2 | 70 kDa | Yes | 3 | 3 |
|  | THAR_000440_T0 | GH31 | 101 kDa | Yes | 4 | 5 |
| Miscellaneous | THAR_000687_T0 | CBM1 | 48 kDa | Yes | 11 | 32 |
|  | THAR_011525_T0 | CBM6 | 50 kDa | Yes | 8 | 15 |
|  | THAR_000186_T0 | CBM20 | 82 kDa | No | 7 | 9 |
|  | THAR_003918_T0 | AA2 | 99 kDa | Yes | 6 | 8 |
|  | THAR_001205_T0 | AA5 | 113 kDa | Yes | 10 | 22 |
|  | THAR_011135_T0 | CE1 | 43 kDa | No | 6 | 17 |
|  | THAR_011592_T0 | CE5 | 24 kDa | Yes | 4 | 12 |
| Chitin | THAR_007593_T0 | GH18 | 46 kDa | Yes | 11 | 33 |

All proteins identified in *T. harzianum* P49P11 secretome is given in supplementary material S4.
<sup>1</sup>Abbreviations: GH, glycoside hydrolase; CBM, carbohydrate- binding module; CE, carbohydrate esterases; PL, polysaccharide lyases; AA, auxiliary activities. CAZymes are represented with the family number according to their representation in the CAZy database.
<sup>2</sup> Prediction of signal peptides based on SignalP analysis.
<sup>3</sup> We only considered proteins identified by at least two unique peptides.
<sup>4</sup> Secretome analysis based on spectral counting.

Oxidoreductases, hemicellulases and beta-glucanases were also detected in P49P11 secretome in response to crystalline cellulose. Among the oxidoreductases, one protein representing AA2, and one AA5 were detected. Although AA2 and AA5 families are peroxidases and copper radical oxidases, respectively, related to modifying lignin [24][25]. strain P49P11 might also use these enzymes in an oxidative-hydrolytic mechanism for cellulose degradation. The presence of hemicellulases and beta-glucanases representing GH11, GH30, GH16, and GH55 suggests enzymes with a broader substrate specificity with weak activities. The secretion of hemicellulases, glucanases, and CAZymes categorized in the miscellaneous group have also been reported in the secretome of *T. reesei, Chaetomium thermophilum*, and *Fusarium metavorans* when grown on different artificial cellulose substrates as the sole carbon source [26][27], suggesting additional CAZymes genes and/or a complex gene regulation for cellulose degradation.

## Conclusion

Here, the cellulolytic potential of *T. harzianum* P49P11 was elucidated using a workflow based on genomic and secretome analyses and enzymatic activities. *T. harzianum* P49P11 genome presented here has a high rate of complete BUSCO genes, which might contribute to a genome-scale understanding of *Trichoderma* species and provide the molecular bases for future omics approaches or genetic engineering. The strain P49P11 genome encodes an arsenal of distinct CAZymes families to degrade different polysaccharides, encouraging new studies to exploit its lignocellulolytic potential. Furthermore, the secretome analysis revealed that strain P49P11 secreted cellulases and non-cellulases in response to crystalline cellulose. This finding suggests a complex gene regulation for cellulose degradation.

## Materials and Methods

### Fungal strain and propagation

The strain *T. harzianum* P49P11 was isolated from Amazon rainforest soil as previous described [9], and deposited at the Embrapa Food Technology Microorganism collection (Rio de Janeiro, Brazil) under accession number BRMCTAA 115. The strain was maintained on potato-dextrose agar (PDA; Difco, BD, USA) medium. For cultivation experiments, agar discs (0.8 cm in diameter) were taken from PDA culture and then transferred to modified Madels medium (MA) [28] containing 1 mL Tween 80, 0.3 g/L urea, 2.0 g/L KH_2_PO_4_, 1.4 g/L (NH_4_)_2_SO_4_, 0.4 g/L CaCl_2_.2H_2_O, 0.3 g/L MgSO_4_.7H_2_O, 5.0 mg/L FeSO4.7H2O, 1.6 mg/L MnSO_4_.4H_2_O, 1.4 mg/L ZnSO_4_.7H_2_O; 2.0 mg/L CoCl_2_.6H_2_O, 1.0 g/L peptone, supplemented with 1 % glucose as carbon source for approximately 3 to 7 days at 29 º C.

### DNA extraction and sequencing

*T. harzianum* P49P11 was culture on PDA plate for 4 days, and then agar discs were transferred to flask containing to modified Madels medium with 1 % glucose. The flask was incubated in shaker on a rotary shaker at 200 rpm of agitation, at 29 º C for 72 hours. The fungal mycelia were collected by filtered, washed three times with distilled water, and then ground into fine powder in liquid nitrogen. The total DNA was extracted using Qiagen DNeasy Plant Mini Kit (QIAGEN) according to the manufacturer’s instructions. Integrity and quality of fungal DNA and the absence of RNA contamination were analyzed using Agilent 2100 Bioanalyzer with 12000 DNA assay kit and agarose gel electrophoresis, respectively (Agilent Technologies, Santa Clara, CA). Total DNA was used to construct Illumina paired-end sequencing library according to the manufacturer’s standard protocol (Illumina, San Diego, CA). Then, library was sequenced on an Illumina HiSeq 2500 system using the paired-end protocol (2×100 bp paired ends) at the Brazilian Biorenewables National Laboratory (LNBR/CNPEM).

### Bioinformatic analysis

Raw sequences quality was checked in FastQC (www.bioinformatics.babraham.ac.uk/projects/fastqc), pre-processed by removing adapters and low-quality sequences using trimmed using Trimmomatic [29] with default setting. Quality-filtered sequences were assembled using Velvet [30], and then the contigs generated were polished and scaffolding using Pilon ([31] and SSPACE [32], respectively. The completeness was estimated with Benchmarking Universal Single-Copy Orthologs (BUSCO) [33]. For genome annotation, gene models were predicted by using the ab initio gene predictors AUGUSTUS [34] and GeneMark-ES [35], and then functional annotation was carried out using InterProScan [36], which used BLASTP search against Gene Ontology (GO) terms [37], COG functional categories[38], Pfam [39], KEGG pathways [40]. Ribosomal RNA (rRNA) and transfer RNA (tRNA) genes were predicted using Infernal [41] and tRNAscan [42], respectively. In addition, genes involved in secondary metabolites were predicted using anitSMASH [43].

For carbohydrate-active enzymes (CAZyme) prediction, all genomic proteins in the *T. harzianum* P49P11 genome were screened for homologs in the dbCAN version 4 database (YIN, et al., 2012) using the HMMER 3.1b package [44] as previously described [45]. To reduce a single proteins sequence with multiple annotations, we manually analyzed the putative CAZymes using PRIAM [46] and Pfam databases, and then best blast hits were used as possible enzymatic activities. Presence of signal peptide in the CAZymes predicted was carried out using SignalP v5 [47].

### Culture condition for enzyme production

For cellulase production and enzyme activities analysis, conidia of *T. harzianum* P49P11 from fresh PDA were harvested and inoculated in 500 mL Erlenmeyer flasks containing 200 mL of Mandels medium supplemented with 1 % glucose as carbon source. The preculture was incubated for 72 h at 29 ° C on a rotary shaker at 250 rpm. The conidia were harvest, washed three times with saline solution, and 10^8^ conidia/mL inoculated into 3.0 L BioFlo 115 fermenter (New Brunswick Scientific Co., USA) with Rushton impeller, with 1.5L final volume. The medium composition in batch fermentation consisted of: modified Mandels medium supplemented with 20 g /L crystalline cellulose (Celufloc®), and foam controlled by polypropylene glycol antifoaming agent (P2000, Dow Chemical, Brazil). The bioreactor was operated under following conditions: agitation 200 - 400 rpm; pH 5.0 - 6.0 controlled by the addition of acid (HCl) or base (NaOH); dissolved O_2_ 30%; temperature 29 °C; air flow 0.35 - 0.7 vvm, minimum oxygen demand set at 30% using agitation and aeration cascade triggered when necessary. Samples of 10 ml were collected at 48 h, 72 h, 96 h and 120 h, and centrifuged at 10,000 rpm for 10 minutes. Supernatants were used to quantify total proteins, and for enzymatic assays and secretome analysis. The experiment was carried out five batch biological replicates.

### Enzymatic activity

Supernatant samples were first filtered through 0.22-µm PVDF (polyvinylidene fluoride) membrane filter and then subjected to enzymatic activities assays. Filter paper hydrolase (FPase), representing total extracellular cellulase activity, was determined according to the methodology described by Ghose (1987), with modifications to reduce the scale of the procedure by a factor of ten. Filter paper activity was assayed by mixing diluted supernatants and sodium acetate buffer (50 mM, pH4.8) with one filter paper strip (Whatman No.1; 1 × 6 cm). The reaction mixture was incubated at 50 ° C for 60 min. For xylanase activity, reaction mixture consisted of 10 μL of the supernatant, 40 μL of 50 mM citrate buffer - pH 4.8 and 50 μL of 0.50% xylan solution (Megazyme, Ireland; Sigma-Aldrich, St. Louis, MO, USA). The reactions were incubated at 50 ° C for 10 minutes. Both reactions were stopped by adding 100 μL of the dinitrosalicylic acid (DNS) reagent followed by heating to 99 °C for 5 minutes. The color change was measured at 540 nm using the Infinite M200® spectrophotometer (Tecan, Switzerland), and total reducing sugar content was estimated by comparison to a glucose standard curve. For β-glycosidase activity, reactions were performed by mixing 10 μL of the supernatant, 40 μL of 50 mM citrate buffer - pH 4.8 and 50 μL of 0.50-nitrophenyl β-D-glucopyranoside 1 mM (pNP-G Sigma) as substrate. Reactions were incubated at 50 ° C for 10 minutes and stopped by adding 100 μL of 1M sodium carbonate (Na_2_CO_3_). The released nitrophenol was quantified by spectrophotometry at 400 nm. All enzymatic measurements were performed in triplicates. One unit of enzyme activity was defined as the amount of enzyme required to release 1 µmol of reducing sugars in 1 min under the assay conditions. Protein concentration in the supernatants was determined according to Bradford (1976).

### Secretome analysis by Liquid Chromatography-Mass Spectrometry

To secretome analysis, crude extracellular supernatant was first subjected to electrophoresis in a denaturing polyacrylamide gel (SDS-PAGE). The gel was stained with Coomassie Brilliant Blue, and the visible bands were cut, reduced, alkylated and submitted to digestion with trypsin according to the internal protocol of LC–MS/MS facilities at National Biosciences Laboratory (LNBio). The peptides obtained were separated by hydrophobicity gradient in a C18 column (75 mm x 100 mm) in an EASY-nLC system chromatography system coupled to an ESI nanospray interface in an LTQ Velos Orbitrap mass spectrometer (Thermo Fisher Scientific). The raw data generated from the mass spectrometry (.raw files) were processed in the Mascot Search Engine program using as parameters a lost cleavage by trypsin, fixed modification of carbamidomethylation, variable modification of methionine oxidation, 10 ppm of mass tolerance for MS, 1 Mass tolerance data for MS / MS. To validate the identification of proteins, the files generated by the Mascot (.dat files) were analyzed by the Scaffold Q + program (Proteome Software Ink., Portland, OR) with parameters for protein identification with a 99% confidence threshold (protein threshold), at least 2 peptides identified for each protein (peptide threshold) and false positive (FDR) equal to or less than 1% and, 50% chance of identifying the amino acid, FDR 0.39%. The data from MS/MS profile was researched and compared against all amino acid sequences of the T. harzianum P49P11.

## Data Availability

The whole genome of *T. harzianum* P49P11 has been deposited in GenBank under Bioproject and Biosample numbers, PRJNA382575 and SAMN06710574, respectively. The secretome data has been deposited in the ProteomeXchange consortium via the PRIDE partner repository [48] under identified PXD033431.

## Supplementary Information

Supplementary Material S1-S4.xlsx

## CRediT authorship contribution statement

C.A.C G.T., and P.S.D: Conceptualized, methodology, investigation, and formal analysis. G.T., and P.S.D: Supervision. G.T: analysed the results and wrote the manuscript – review & editing. G.T., G.F.P., and D.M.RP. performed the bioinformatic analyses. C.A.C., P.S.D., G.F.P., D.M.RP., F.M.S., J.G.C.P revised the manuscript. All authors read and approved the final manuscript.

## Acknowledgements

We gratefully acknowledge the provision of time on the MAS facilities (LNBio) at the National Center for Research in Energy and Materials (CNPEM) and Brazilian Biorenewables National Laboratory (LNBr) NGS Sequencing Facility for generating the sequencing data described here.

## Reference

[1] Drula E, Garron M-L, Dogan S, Lombard V, Henrissat B, Terrapon N. The carbohydrate-active enzyme database: functions and literature. Nucleic Acids Res 2021:1–7. https://doi.org/10.1093/nar/gkab1045.

[2] Grand View Research. Enzymes Market 2021:104.

[3] Paul M, Mohapatra S, Kumar Das Mohapatra P, Thatoi H. Microbial cellulases – An update towards its surface chemistry, genetic engineering and recovery for its biotechnological potential. Bioresour Technol 2021;340. https://doi.org/10.1016/j.biortech.2021.125710.

[4] Saini S, Sharma KK. Fungal lignocellulolytic enzymes and lignocellulose: A critical review on their contribution to multiproduct biorefinery and global biofuel research. Int J Biol Macromol 2021;193:2304–19. https://doi.org/10.1016/j.ijbiomac.2021.11.063.

[5] Madhavan A, Arun KB, Sindhu R, Alphonsa Jose A, Pugazhendhi A, Binod P, et al. Engineering interventions in industrial filamentous fungal cell factories for biomass valorization. Bioresour Technol 2022;344:126209. https://doi.org/10.1016/j.biortech.2021.126209.

[6] Amini Z, Self R, Strong J, Speight R, O’Hara I, Harrison MD. Valorization of sugarcane biorefinery residues using fungal biocatalysis. Biomass Convers Biorefinery 2022;12:997–1011. https://doi.org/10.1007/s13399-021-01456-3.

[7] Cherry JR, Fidantsef AL. Directed evolution of industrial enzymes: An update. Curr Opin Biotechnol 2003;14:438–43. https://doi.org/10.1016/S0958-1669(03)00099-5.

[8] Uusitalo JM, Helena Nevalainen KM, Harkki AM, Knowles JKC, Penttilä ME. Enzyme production by recombinant Trichoderma reesei strains. J Biotechnol 1991;17:35–49. https://doi.org/10.1016/0168-1656(91)90025-Q.

[9] Delabona P da S, Farinas CS, da Silva MR, Azzoni SF, Pradella JG da C. Use of a new Trichoderma harzianum strain isolated from the Amazon rainforest with pretreated sugar cane bagasse for on-site cellulase production. Bioresour Technol 2012;107:517–21. https://doi.org/10.1016/j.biortech.2011.12.048.

[10] Delabona P da S, Lima DJ, Robl D, Rabelo SC, Farinas CS, da Cruz Pradella JG. Enhanced cellulase production by Trichoderma harzianum by cultivation on glycerol followed by induction on cellulosic substrates. J Ind Microbiol Biotechnol 2016;43:617–26. https://doi.org/10.1007/s10295-016-1744-8.

[11] Souza MF de, Silva ASA da, Bon EPS. A novel Trichoderma harzianum strain from the Amazon Forest with high cellulolytic capacity. Biocatal Agric Biotechnol 2018;14:183–8. https://doi.org/10.1016/j.bcab.2018.03.008.

[12] Li JX, Zhang F, Jiang DD, Li J, Wang F Lou, Zhang Z, et al. Diversity of Cellulase-Producing Filamentous Fungi From Tibet and Transcriptomic Analysis of a Superior Cellulase Producer Trichoderma harzianum LZ117. Front Microbiol 2020;11:1–15. https://doi.org/10.3389/fmicb.2020.01617.

[13] Almeida DA, Horta MAC, Ferreira Filho JA, Murad NF, de Souza AP. The synergistic actions of hydrolytic genes reveal the mechanism of Trichoderma harzianum for cellulose degradation. J Biotechnol 2021;334:1–10. https://doi.org/10.1016/j.jbiotec.2021.05.001.

[14] Gelain L, Kingma E, Geraldo da Cruz Pradella J, Carvalho da Costa A, van der Wielen L, van Gulik WM. Continuous production of enzymes under carbon-limited conditions by Trichoderma harzianum P49P11. Fungal Biol 2021;125:177–83. https://doi.org/10.1016/j.funbio.2020.10.008.

[15] Delabona P da S, Cota J, Hoffmam ZB, Paixão DAA, Farinas CS, Cairo JPLF, et al. Understanding the cellulolytic system of Trichoderma harzianum P49P11 and enhancing saccharification of pretreated sugarcane bagasse by supplementation with pectinase and α-l-arabinofuranosidase. Bioresour Technol 2013;131:500–7. https://doi.org/10.1016/j.biortech.2012.12.105.

[16] Druzhinina IS, Kubicek CP. Genetic engineering of Trichoderma reesei cellulases and their production. Microb Biotechnol 2017. https://doi.org/10.1111/1751-7915.12726.

[17] Yan S, Xu Y, Yu X-W. From induction to secretion: a complicated route for cellulase production in Trichoderma reesei. Bioresour Bioprocess 2021;8. https://doi.org/10.1186/s40643-021-00461-8.

[18] Kubicek CP, Steindorff AS, Chenthamara K, Manganiello G, Henrissat B, Zhang J, et al. Evolution and comparative genomics of the most common Trichoderma species. BMC Genomics 2019;20:1–24. https://doi.org/10.1186/s12864-019-5680-7.

[19] Kubicek CP, Herrera-Estrella A, Seidl-Seiboth V, Martinez DA, Druzhinina IS, Thon M, et al. Comparative genome sequence analysis underscores mycoparasitism as the ancestral life style of Trichoderma. Genome Biol 2011;12. https://doi.org/10.1186/gb-2011-12-4-r40.

[20] Rytioja J, Hildén K, Yuzon J, Hatakka A, de Vries RP, Mäkelä MR. Plant-Polysaccharide-Degrading Enzymes from Basidiomycetes. Microbiol Mol Biol Rev 2014;78:614–49. https://doi.org/10.1128/MMBR.00035-14.

[21] Ferreira Filho JA, Horta MAC, dos Santos CA, Almeida DA, Murad NF, Mendes JS, et al. “Integrative genomic analysis of the bioprospection of regulators and accessory enzymes associated with cellulose degradation in a filamentous fungus (Trichoderma harzianum).” BMC Genomics 2020;21:1–14. https://doi.org/10.1186/s12864-020-07158-w.

[22] Zhu N, Yang J, Ji L, Liu J, Yang Y, Yuan H. Metagenomic and metaproteomic analyses of a corn stover-adapted microbial consortium EMSD5 reveal its taxonomic and enzymatic basis for degrading lignocellulose. Biotechnol Biofuels 2016;9:243. https://doi.org/10.1186/s13068-016-0658-z.

[23] Paixao DAA, Tomazetto G, Sodre VR, Gonçalves TA, Uchima CA, Büchli F, et al. Microbial enrichment and meta-omics analysis identify CAZymes from mangrove sediments with unique properties. Enzyme Microb Technol 2021;148:1–12. https://doi.org/10.1016/j.enzmictec.2021.109820.

[24] Levasseur A, Drula E, Lombard V, Coutinho PM, Henrissat B. Expansion of the enzymatic repertoire of the CAZy database to integrate auxiliary redox enzymes. Biotechnol Biofuels 2013;6:41. https://doi.org/10.1186/1754-6834-6-41.

[25] Daou M, Faulds CB. Glyoxal oxidases: their nature and properties. World J Microbiol Biotechnol 2017;33:1–11. https://doi.org/10.1007/s11274-017-2254-1.

[26] Jiang X, Du J, He R, Zhang Z, Qi F, Huang J, et al. Improved Production of Majority Cellulases in Trichoderma reesei by Integration of cbh1 Gene From Chaetomium thermophilum. Front Microbiol 2020;11. https://doi.org/10.3389/fmicb.2020.01633.

[27] Brandt SC, Brognaro H, Ali A, Ellinger B, Maibach K, Rühl M, et al. Insights into the genome and secretome of Fusarium metavorans DSM105788 by cultivation on agro-residual biomass and synthetic nutrient sources. Biotechnol Biofuels 2021;14:1–22. https://doi.org/10.1186/s13068-021-01927-9.

[28] Mandels M, Reese ET. Induction of cellulase in Trichoderma viride as influenced by carbon source and metals 1956:269–78.

[29] Bolger AM, Lohse M, Usadel B. Trimmomatic: A flexible trimmer for Illumina sequence data. Bioinformatics 2014;30:2114–20. https://doi.org/10.1093/bioinformatics/btu170.

[30] Zerbino DR, Birney E. Velvet: Algorithms for de novo short read assembly using de Bruijn graphs. Genome Res 2008;18:821–9. https://doi.org/10.1101/gr.074492.107.

[31] Walker BJ, Abeel T, Shea T, Priest M, Abouelliel A, Sakthikumar S, et al. Pilon: An integrated tool for comprehensive microbial variant detection and genome assembly improvement. PLoS One 2014;9. https://doi.org/10.1371/journal.pone.0112963.

[32] Boetzer M, Henkel C V, Jansen HJ, Butler D, Pirovano W. Scaffolding pre-assembled contigs using SSPACE. Bioinformatics 2011;27:578–9.

[33] Manni M, Berkeley MR, Seppey M, Simao FA, Zdobnov EM. BUSCO update : novel and streamlined workflows along with broader and deeper. Mol Biol Evol 2021. https://doi.org/https://doi.org/10.1093/molbev/msab199.

[34] Stanke M, Diekhans M, Baertsch R, Haussler D. Using native and syntenically mapped cDNA alignments to improve de novo gene finding. Bioinformatics 2008;24:637–44. https://doi.org/10.1093/bioinformatics/btn013.

[35] Ter-Hovhannisyan V, Lomsadze A, Chernoff YO, Borodovsky M. Gene prediction in novel fungal genomes using an ab initio algorithm with unsupervised training. Genome Res 2008;18:1979–90. https://doi.org/10.1101/gr.081612.108.

[36] Blum M, Chang HY, Chuguransky S, Grego T, Kandasaamy S, Mitchell A, et al. The InterPro protein families and domains database: 20 years on. Nucleic Acids Res 2021;49:D344–54. https://doi.org/10.1093/nar/gkaa977.

[37] Carbon S, Douglass E, Good BM, Unni DR, Harris NL, Mungall CJ, et al. The Gene Ontology resource: Enriching a GOld mine. Nucleic Acids Res 2021;49:D325–34. https://doi.org/10.1093/nar/gkaa1113.

[38] Galperin MY, Wolf YI, Makarova KS, Alvarez RV, Landsman D, Koonin E V. COG database update: Focus on microbial diversity, model organisms, and widespread pathogens. Nucleic Acids Res 2021;49:D274–81. https://doi.org/10.1093/nar/gkaa1018.

[39] Mistry J, Chuguransky S, Williams L, Qureshi M, Salazar GA, Sonnhammer ELL, et al. Pfam: The protein families database in 2021. Nucleic Acids Res 2021;49:D412–9. https://doi.org/10.1093/nar/gkaa913.

[40] Kanehisa M, Goto S, Sato Y, Furumichi M, Tanabe M. KEGG for integration and interpretation of large-scale molecular data sets. Nucleic Acids Res 2012;40:D109--14. https://doi.org/10.1093/nar/gkr988.

[41] Nawrocki EP, Eddy SR. Infernal 1.1: 100-fold faster RNA homology searches. Bioinformatics 2013;29:2933–5. https://doi.org/10.1093/bioinformatics/btt509.

[42] Lowe TM, Eddy SR. tRNAscan-SE: A program for inproved detection of transfer RNA genes in genomic sequence. Nucleic Acids Res 1997;25:955–64. https://doi.org/10.1093/nar/25.5.0955.

[43] Blin K, Shaw S, Kloosterman AM, Charlop-Powers Z, Van Wezel GP, Medema MH, et al. AntiSMASH 6.0: Improving cluster detection and comparison capabilities. Nucleic Acids Res 2021;49:W29–35. https://doi.org/10.1093/nar/gkab335.

[44] Eddy SR. Accelerated profile HMM searches. PLoS Comput Biol 2011;7. https://doi.org/10.1371/journal.pcbi.1002195.

[45] Tomazetto G, Pimentel AC, Wibberg D, Dixon N, Squina FM. Multi-omic Directed Discovery of Cellulosomes, Polysaccharide Utilization Loci, and Lignocellulases from an Enriched Rumen Anaerobic Consortium. Appl Environ Microbiol 2020;86:1–23. https://doi.org/https://doi.org/10.1128/AEM.00199-20.

[46] Claudel-Renard C, Chevalet C, Faraut T, Kahn D. Enzyme-specific profiles for genome annotation: PRIAM. Nucleic Acids Res 2003;31:6633–9. https://doi.org/10.1093/nar/gkg847.

[47] Almagro Armenteros JJ, Tsirigos KD, Sønderby CK, Petersen TN, Winther O, Brunak S, et al. SignalP 5.0 improves signal peptide predictions using deep neural networks. Nat Biotechnol 2019;37:420–3. https://doi.org/10.1038/s41587-019-0036-z.

[48] Perez-Riverol Y, Csordas A, Bai J, Bernal-Llinares M, Hewapathirana S, Kundu DJ, et al. The PRIDE database and related tools and resources in 2019: Improving support for quantification data. Nucleic Acids Res 2019;47:D442–50. https://doi.org/10.1093/nar/gky1106.

